# Cofilin pathology is a new player on α-synuclein-induced spine impairment in models of hippocampal synucleinopathy

**DOI:** 10.1101/2021.02.02.425931

**Authors:** MI Oliveira da Silva, M Santejo, IW Babcock, A Magalhães, LS Minamide, E Castillo, E Gerhardt, C Fahlbusch, RA Swanson, TF Outeiro, JR Bamburg, MA Liz

## Abstract

Cognitive dysfunction and dementia are presently recognized as major complications in α-synucleinopathies, namely in Dementia with Lewy Bodies (DLB) and Parkinson’s disease with dementia (PDD). In these disorders, α-Synuclein (αSyn) accumulation affects severely the hippocampus by inducing synaptic dysfunction which culminates in cognitive impairment. To characterize the mechanisms underlying αSyn-induced neuronal dysfunction we analysed the effect of overexpression or extracellular administration of αSyn on hippocampal neurons. We observed that αSyn induces the dysregulation of the actin-binding protein cofilin and its assembly into rod structures in a mechanism mediated by the cellular prion protein (PrP^C^). Moreover, we unraveled cofilin pathology as mediator of αSyn-induced dendritic spine impairment in hippocampal neurons. Importantly, in a synucleinopathy mouse model with cognitive impairment we validated cofilin dysregulation and synaptic dysfunction at the same age when cognitive deficits were observed. Our data supports cofilin as a novel player on hippocampal synaptic dysfunction triggered by αSyn on Lewy Body dementias.

## Introduction

α-Synuclein (αSyn) is well-known due to its involvement in Parkinson’s disease (PD), a neurodegenerative disorder mainly characterized by motor dysfunction^1^, in which the major hallmarks are the formation of neuronal αSyn-containing inclusions, known as Lewy Bodies, and the loss of dopaminergic neurons in the substantia nigra pars compacta (SNpc)^2, 3^. Nevertheless, αSyn is implicated in other synucleinopathies namely Lewy Body dementias, which include Dementia with Lewy Bodies (DLB) and Parkinson’s disease with dementia (PDD). These two disorders present similar symptomatology, being mainly distinguishable by the time at which dementia occurs. In DLB, cognitive impairment precedes parkinsonism or begins within a year of parkinsonism, whereas in PDD, parkinsonism generally precedes cognitive impairment by more than 1 year^4^. DLB accounts for ∼30% of all age-related dementias. In PD, 20–40% of the patients have cognitive impairments at disease onset, and ∼80% of the patients develop PDD^5^. Despite the impact on patients, no treatments have been proven to slow or stop disease progression in Lewy Body dementias, and the pathophysiological mechanisms underlying these disorders require further investigation.

Lewy Body dementias have been linked to increased expression/aggregation of αSyn in the hippocampal brain region leading to dysfunction. Locus multiplications of the *SNCA* gene have been described in PDD and DLB cases, whose severity of cognitive impairment and age of onset correlate with the copy number of the *SNCA* gene^6^. The hippocampus, a major brain component with key roles in memory and learning, is one of the most vulnerable regions affected by αSyn pathology^4^. In this respect, hippocampal volume loss is observed in DLB and PDD patients, but not in cases of PD with normal cognition^7^. Moreover, increased levels of Lewy Body pathology are observed in the hippocampus of Lewy Body dementias^8^. These observations suggest that the impact of αSyn on the hippocampus may underlie cognitive deficits and dementia observed in Lewy Body dementias. Additionally, synaptic dysfunction in the hippocampus of mouse models of synucleinopathies evidence the link between cognitive deficits and hippocampal Lewy Body pathology.

In this study, we aimed at characterizing mechanisms downstream of αSyn hippocampal pathology underlying synaptic impairment and cognitive dysfunction in Lewy Body dementias. We focused on the actin cytoskeleton, not only due to its critical role in synaptic function, but also because a link between this cell component and αSyn has been increasingly supported by the literature^9, 10^. One of the consequences of the dysregulation of actin dynamics in neurodegenerative disorders is the formation of cofilin-actin rods. These are structures composed by bundles of cofilin-saturated actin filaments, which result from the dysregulation of the actin binding protein cofilin *via* localized hyperactivation by dephosphorylation and oxidation, and that were shown to block intracellular trafficking and induce synaptic loss in hippocampal neurons^11^. Cofilin-actin rods have been mainly implicated in cognitive impairment in Alzheimer’s disease (AD)^12, 13^ and synaptic dysfunction after ischemic stroke^14, 15^. Importantly, decreasing either cofilin dephosphorylation or the total levels of cofilin expression was effective in the amelioration of cofilin pathology and cognitive deficits in the Aβ-overproducing mouse model of AD^16-18^. Additionally, in a rat model of ischemic stroke, which presents rod formation in the peri-infarct area, the overexpression of active LIM kinase (LIMK), which phosphorylates cofilin at Ser3, promoting its inactivation, resulted in decreased rod formation and rescue of synaptic activity^14^. These data suggest that modulation of cofilin pathology is beneficial to preserve synaptic function and prevent cognitive deficits in these neurological disorders.

Here we show that αSyn induces cofilin pathology in hippocampal neurons *via* a cellular prion protein (PrP^C^)-dependent pathway. Moreover, cofilin dysregulation mediates dendritic spine impairment in response to αSyn. Based on the present data, we propose that αSyn-induced cofilin pathology may underlie synaptic dysfunction and cognitive impairment in Lewy Body dementias.

## Results

### αSyn induces cofilin-actin rod formation in hippocampal neurons *via* PrP^c^

In order to characterize downstream mechanisms underlying hippocampal dysfunction in Lewy Body dementias, we analysed the effect of overexpressing wild type (WT) αSyn in primary cultures of hippocampal neurons to establish a scenario of increased levels and aggregation of αSyn^19^which were shown to induce synaptic dysfunction^20^. Rat hippocampal neurons were infected at DIV4 with lentivirus encoding for WT αSyn-IRES-GFP or IRES-GFP as control. Overexpression of αSyn, which was highly phosphorylated at Ser129, was confirmed by western blot and immunostaining of DIV7 transduced neurons (Figure 1A and B). Similar analyses at DIV14, a time point when endogenous αSyn is already expressed and enriched in presynaptic terminals, allowed a quantitative analysis of αSyn overexpression relatively to the levels of endogenous protein. We observed that in neurons infected at DIV4 with WT αSyn-IRES-GFP lentivirus, and analyzed at DIV14, αSyn levels were increased approximately 3-fold when compared to control cells, with similar levels of GFP expression (Figure S1A and B). Immunocytochemistry at DIV14 revealed the presence of endogenous αSyn, without detection of αSyn pS129, in control cells (Figure S1C). This result is in accordance with the phosphorylation levels of αSyn pS129 in healthy brains, which correspond to about 4% of the total protein^21^. In the case of WT αSyn-transduced neurons, αSyn was extensively phosphorylated at Ser129, a marker for aggregation of the protein, mimicking the aberrant accumulation of αSyn pS129 in the brain of patients with Lewy Body disease^22^ (Figure S1C). These results confirm that we have successfully established a cell system of αSyn hippocampal pathology.

**Figure 1.**
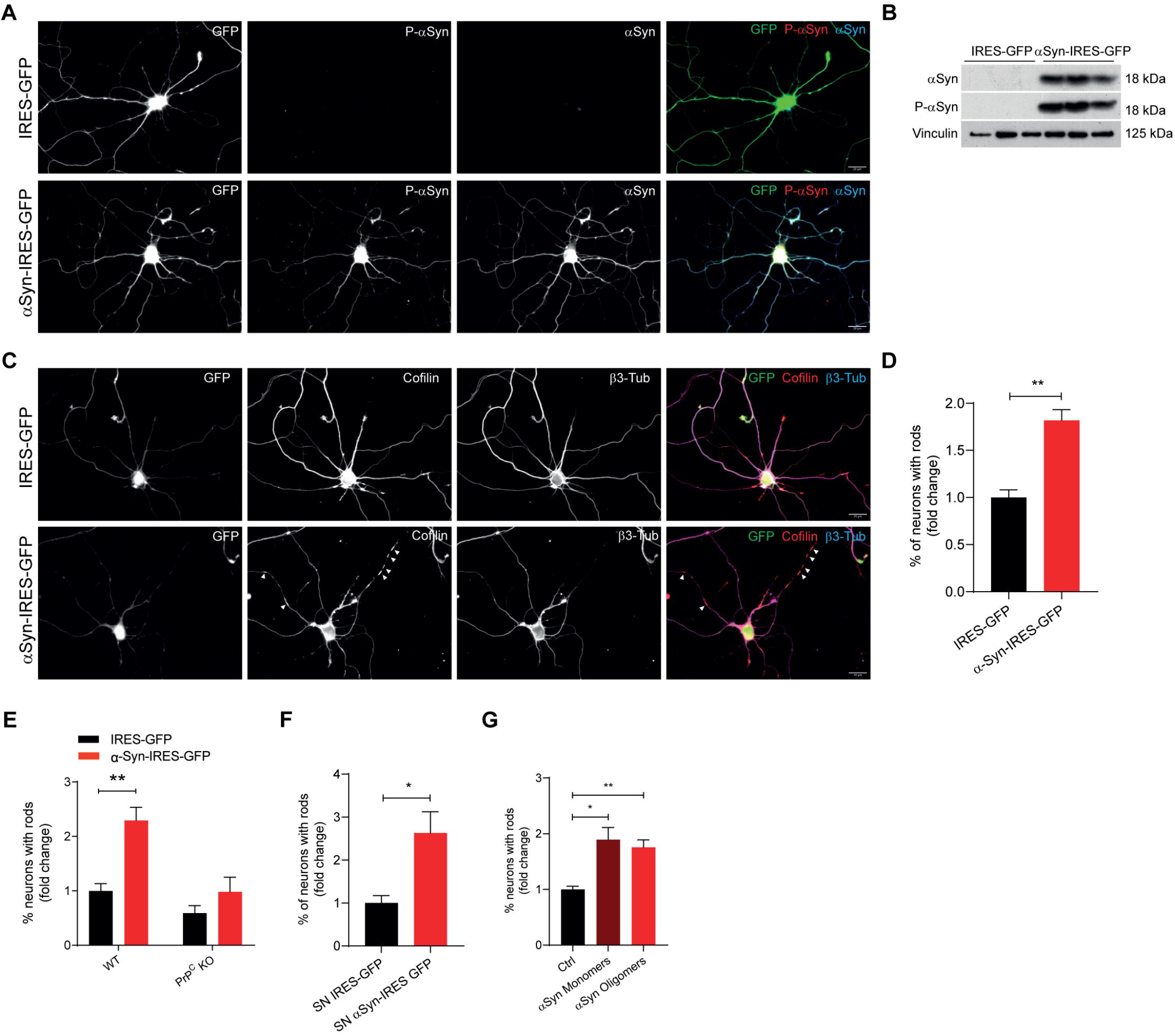
αSyn induces cofilin-actin rods formation in hippocampal neurons via PrP^C^. **(A-D)** Primary hippocampal neurons were infected at DIV4 with IRES-GFP or WT-αSyn-IRES-GFP lentivirus and analyzed at DIV7. **(A)** Representative images of infected hippocampal neurons immunostained with α-Syn pS129 (red) and α-Syn (blue). Scale bar: 20μm. **(B)** Western blot analysis of α-Syn and α-Syn pS129 levels in transduced hippocampal neurons. Vinculin was used as loading control. Data represent mean±SEM (n=3 replicates/treatment). **(C)** Representative images of infected hippocampal neurons immunostained with Cofilin (red) and β3-Tubulin (blue). Scale bar: 20μm. Arrowheads indicate cofilin-actin rod structures. **(D)** Percentage of neurons with rods relative to **C**. Data represent mean±SEM (n=3 replicates/treatment with ≥100 neurons/replicate). **p<0.01 by Student’s *t* test. **(E)** Primary hippocampal neurons from WT or PrP^C^ KO mice were infected at DIV4 with IRES-GFP or WT-αSyn-IRES-GFP lentivirus and the percentage of neurons with rods (shown as fold change relative to control) was analyzed at DIV7. Data represent mean±SEM (n=5-6 replicates/treatment with ≥100 neurons/replicate). **p<0.01 by Student’s *t* test. **(F)** Percentage of neurons with rods (shown as fold change relative to control) on DIV7 hippocampal neurons previously treated for 48h with supernatants from IRES-GFP or WT-αSyn-IRES-GFP infected neurons. Data represent mean±SEM (n=4 replicates/treatment with ≥100 neurons/condition). *p<0.05 by Student’s *t* test. **(G)** Percentage of neurons with rods on DIV7 hippocampal neurons pre-treated with Control (PBS), αSyn monomers (0.5μM) or αSyn oligomers (0.5μM) for 24h. Data represent mean±SEM (n=3 replicates/treatment with ≥100 neurons/condition). *p<0.05, **p<0.01 by Student’s *t* test.

Cofilin-actin rods are one of the features associated with hippocampal pathology in response to Aβ, leading to synaptic impairment and cognitive dysfunction in AD^16^. We hypothesized whether αSyn would exert a similar effect in the context of hippocampal pathology. Using our established cell system of αSyn hippocampal pathology at DIV7, we demonstrated that αSyn overexpression induced a significant 1.8-fold increase in the percentage of neurons presenting rod formation when compared to control neurons (Figure 1C and D). Interestingly, the percentage of neurons in WT αSyn-transduced neurons was approximately of 23%, a similar value to the one reported for Aβ-induced rod formation in rat hippocampal neurons^23, 24^. Moreover, in AD, Aβ-induced formation of rods occurs *via* a PrP^C^-dependent pathway leading to NADPH oxidase (NOX) activation^25^, which culminates in the localized dysregulation of cofilin activity *via* oxidation and dephosphorylation. Thus, we addressed whether, similarly to Aβ, αSyn could act *via* PrP^C^. In order to test this hypothesis, we overexpressed αSyn using the established lentiviral strategy in hippocampal neurons from either WT or PrP^C^ knockout (KO) mice. αSyn overexpression induced rods in neurons from WT mice, leading to a 2.3-fold increase in the percentage of neurons forming rods when compared to control transduced neurons. In the case of PrP^C^ KO neurons, αSyn was not able to induce rods formation (Figure 1E).

Our data confirm a common molecular mechanism for rod formation between αSyn and Aβ involving PrP^C^. PrP^C^ is a lipid-linked protein on the membrane outer surface that was shown to be important for the transfer of αSyn between cells leading to spreading of pathology^26^. More recently, it was demonstrated that PrP^C^ interaction with αSyn existent in extracellular milieu, triggers a signaling pathway that culminates in synaptic dysfunction in hippocampal neurons^27^. Based on that observation, and on the fact that αSyn was shown to be released into the culture medium in neurons overexpressing the protein, we analyzed whether αSyn released from neurons had an impact in rod formation. Dot blot analysis performed with nitrocellulose membrane confirmed the presence of exogenous αSyn in the culture media collected from αSyn-transduced hippocampal neurons (Figure S2). Additional analysis with a cellulose acetate membrane, which retain protein aggregates, did not detect αSyn aggregated species (data not shown), suggesting that the culture media of αSyn transduced hippocampal neurons contains soluble monomers/oligomers of αSyn. Addition of the αSyn-containing media for 48h to DIV5 wild-type hippocampal neurons, which have no detectable levels of endogenous of αSyn, induced a significant 2.6-fold increase in the percentage of neurons with rods, when compared to cells treated with control media, with approximately 20% of the neurons presenting rod formation (Figure 1F). Interestingly, the exogenous addition of recombinant αSyn monomeric and oligomeric species had a similar effect in rod formation (Figure 1G). These results show that αSyn-overexpressing hippocampal neurons release soluble αSyn that induces cofilin-actin rods in non-αSyn-transduced neurons. Although these data might suggest that the presence of αSyn in the extracellular milieu is associated with induction of rod formation *via* PrP^C^, we cannot discard an intracellular effect of the overexpressed and aggregated αSyn.

### Cofilin-actin rod formation is recapitulated in a model of αSyn PFFs-induced pathology

The exogenous addition of αSyn preformed fibrils (PFFs), derived from the aggregation of recombinant αSyn monomers, was shown to promote formation of αSyn pathogenic inclusions in WT neurons and mice, as well as being involved in αSyn spreading and synaptic dysfunction in hippocampal neurons^28^. Therefore, we aimed to also analyze rod formation in a model of exogenous addition of αSyn PFFs. Initially, and considering an extracellular effect of the exogenous addition of αSyn on rod formation, we questioned whether a 24h addition of 1 µg/mL of αSyn PFFs to DIV7 hippocampal neurons could induce rod formation. We observed that treatment of WT mouse hippocampal neurons induced a 3.1-fold increase in the percentage of neurons with rods (Figure 2A). Similarly to what we observed in αSyn-overexpressing neurons, the αSyn PFFs effect on rod formation was abolished when using PrP^C^KO neurons (Figure 2A). Additionally, to confirm the involvement of NOX in αSyn induced rod formation, we used neurons from p47 KO mice and observed that αSyn PFFs were not able to promote rod formation (Figure 2A). This data suggests an extracellular effect of αSyn PFFs on rod induction, which seems not to be related with seeding and intracellular aggregation, as on one hand, DIV7 neurons do not show detectable levels of endogenous protein (Figure 1A), and on the other hand, immunocytochemistry analysis of DIV7 neurons pre-treated with αSyn PFFs did not detect αSyn pS129 (data not shown).

**Figure 2.**
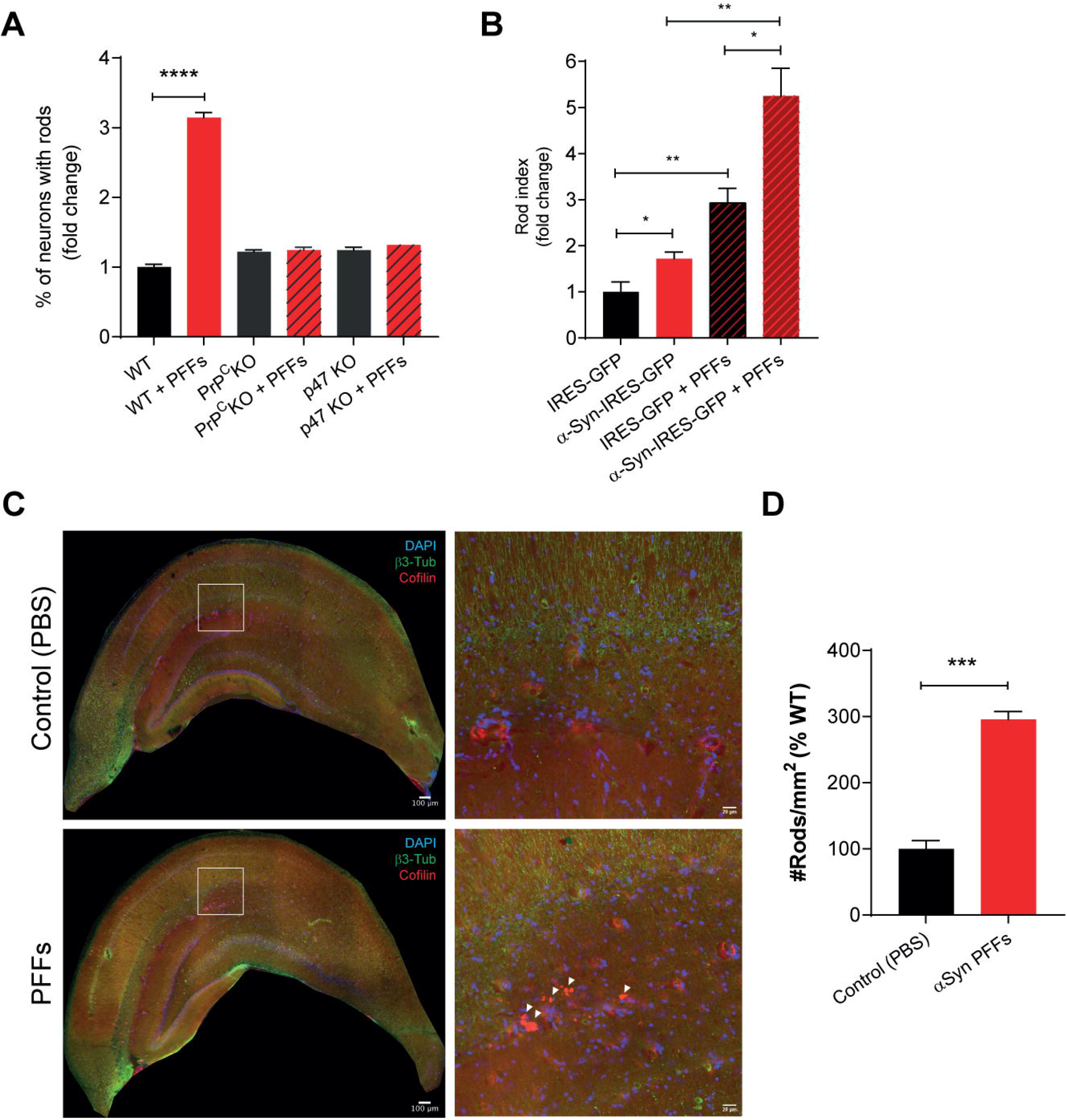
αSyn pre-formed fibrils induce cofilin-actin rods formation in hippocampal neurons. **(A)** Primary hippocampal neurons from WT, PrP^C^ KO or p47 KO mice treated with control (PBS) or αSyn PFFs 1μg/ml at DIV6 and the percentage of neurons with rods (shown as fold change relative to control) was analyzed at DIV7. Data represent mean±SEM (n=3 replicates/treatment with ≥100 neurons/replicate). ****p<0.0001 by Student’s *t* test. **(B)** Rod index (Number of rods/number of neurons, shown as fold change relative to control) of hippocampal neurons infected with IRES-GFP or WT-αSyn-IRES-GFP lentivirus at DIV4 with or without addition of αSyn PFFs (150µg/mL) at DIV7 and analyzed at DIV14. Data represent mean±SEM (n=3 replicates/treatment with ≥100 neurons/condition). *p<0.05, **p<0.01 by Student’s *t* test. **(C)** Representative images of brain sections from Control (PBS) and αSyn PFFs injected mice 7 months after the injection. Sections stained for DAPI (blue), β3-Tubulin (green) and Cofilin (red) and the respective insets in the hippocampal region. Scale bar: 100μm and 20μm for insets. Arrowheads indicate cofilin-actin rod structures. **(D)** Number of rods per mm^2^ (percentage relative to WT) in the hippocampal region relative to **C**. Data represent mean±SEM (n=3-4 animals/condition with 3 sections/animal). ***p<0.001 by Student’s t test.

To analyse rod formation induced by αSyn PFFs in a more physiologic context we performed analysis of DIV14 neurons, when endogenous αSyn is detectable and synapses are mature. In this experiment hippocampal neurons were infected at DIV4 with either WT αSyn-IRES-GFP or IRES-GFP lentivirus, pre-treated or not with 150ng/mL of αSyn PFFs at DIV7, and analyzed for rod formation at DIV14. As seen previously at DIV7, αSyn-overexpression induced rod formation in mature neurons as observed by the 1.7-fold increase in rod index (Figure 2B). αSyn PFFs induced a 2.9-fold increase in rod index in GFP-transduced cells while in αSyn-transduced neurons the increment in rod formation was of approximately 3-fold (when compared to αSyn-transduced neurons) (Figure 2B). In this experiment we detected by immunocytochemistry high levels of αSyn pS129 in WT αSyn-transduced neurons (Figure S1C) and in GFP-transduced neurons treated with αSyn PFFs, being the highest levels of αSyn pS129 observed in WT αSyn-transduced neurons with αSyn PFFs (data not shown). Altogether, our findings suggest that the process of spreading, seeding and intracellular accumulation of αSyn pS129 contributes, but seems not to be required (Figure 2A), for αSyn-induced rod formation.

Aiming at assessing whether αSyn PFFs have an effect on rod formation *in vivo*, we used a mouse model of αSyn PFFs injection in the substantia nigra. This model was chosen to analyze the effect of spreading *in vivo*, as αSyn species injected in the substantia nigra pars compacta were found in the cortex, striatum, thalamus, and hippocampus^26^. We observed the formation of cofilin-actin rods in the hippocampus of mice 7 months post-injection, which presents a 2.9-fold increase in rod formation compared to control injected mice (Figure 2C and D). These results recapitulate our *in vitro* data of αSyn PFFs-induced rod formation in hippocampal neurons. More importantly, these findings further support that spreading of αSyn species contributes to cofilin pathology.

### Cofilin activation mediates αSyn-induced rod formation and dendritic spine impairment

Although we demonstrated rod formation in hippocampal neurons in different scenarios of αSyn-induced pathology, we followed our *in vitro* experiments in the context of αSyn-overexpression in DIV14 mature neurons. This was due to our ultimate goal which was to validate our findings *in vivo* using a mouse model of neuronal αSyn-overexpression as shown in the following section (Figure S3).

Cofilin-actin rod formation results from the local activation of cofilin by dephosphorylation at the Ser3 residue. To test the impact of αSyn on the activation status of cofilin, we performed western blot analysis of cofilin pS3 and total cofilin on cell extracts from DIV14 cultured hippocampal neurons infected either with WT αSyn-IRES-GFP or IRES-GFP lentivirus. We observed that αSyn overexpression induced a significant decrease in the levels of cofilin pS3 with no alterations in the levels of total cofilin (Figure 3A and B). Taking into consideration that αSyn induced rods in approximately 20% of the transduced neurons, it was surprising that we detected a global decrease in cofilin phosphorylation by western blot. In the case of Aβ treatment, changes in cofilin phosphorylation were not detected by western blot as those were shown to occur only in rod-forming neurites^25^. Our data demonstrates that αSyn induces a global cofilin activation by dephosphorylation in hippocampal neurons. Having established evidence for cofilin activation in the αSyn overexpressing neurons, and since cofilin has key roles in regulating actin dynamics in neurons, we assessed the impact of cofilin activation induced by αSyn on hippocampal neuronal function. Initially, we evaluated rod formation in DIV14 neurons transfected at DIV12 with the lentiviral plasmids expressing either WT αSyn-IRES-GFP or IRES-GFP. We determined a 1.8-fold increase in rod index in cells transfected with the WT αSyn-IRES-GFP plasmid when compared to cells transfected with IRES-GFP control plasmid (Figure 3C and D). To validate that αSyn-induced rod formation was a result of cofilin activation, we performed co-transfection with a cofilin phospho-mimetic inactive mutant (cofilin-S3E). Whereas cofilin-S3E had no effect on rod index in control cells, expression of the mutant totally abolished αSyn-induced rod formation (Figure 3C and D), confirming that cofilin activation induced by αSyn mediates rod formation. To further characterize the impact of αSyn-induced cofilin pathology on neurons, we assessed neuronal function focusing on dendritic spine density and morphology. Dendritic spines are small protrusions from the dendritic shaft and their proper regulation is critical, with cofilin being one of the key players in this process, as its tight regulation is crucial for spine formation and maintenance^29^. Using our neuronal culture system (Figure 3C and D), we found that αSyn-expressing neurons presented a significant decrease in dendritic spine number when compared with control neurons (Figure 3E and F). The observed reduction in dendritic spines resulted mainly from a decrease in mushroom (mature) and filopodium (immature) spine types (Figure 3E and G). This suggests that αSyn and consequent cofilin pathology affects not only spine maturation and maintenance but can also impact in stages of spine formation. Additionally, the alterations in spine density and morphology induced by αSyn were significantly reverted, although not to control levels, when cofilin S3E was co-expressed with αSyn (Figure 3E-G), unraveling cofilin activation as a novel player on αSyn-induced dendritic spine impairment in hippocampal neurons.

**Figure 3.**
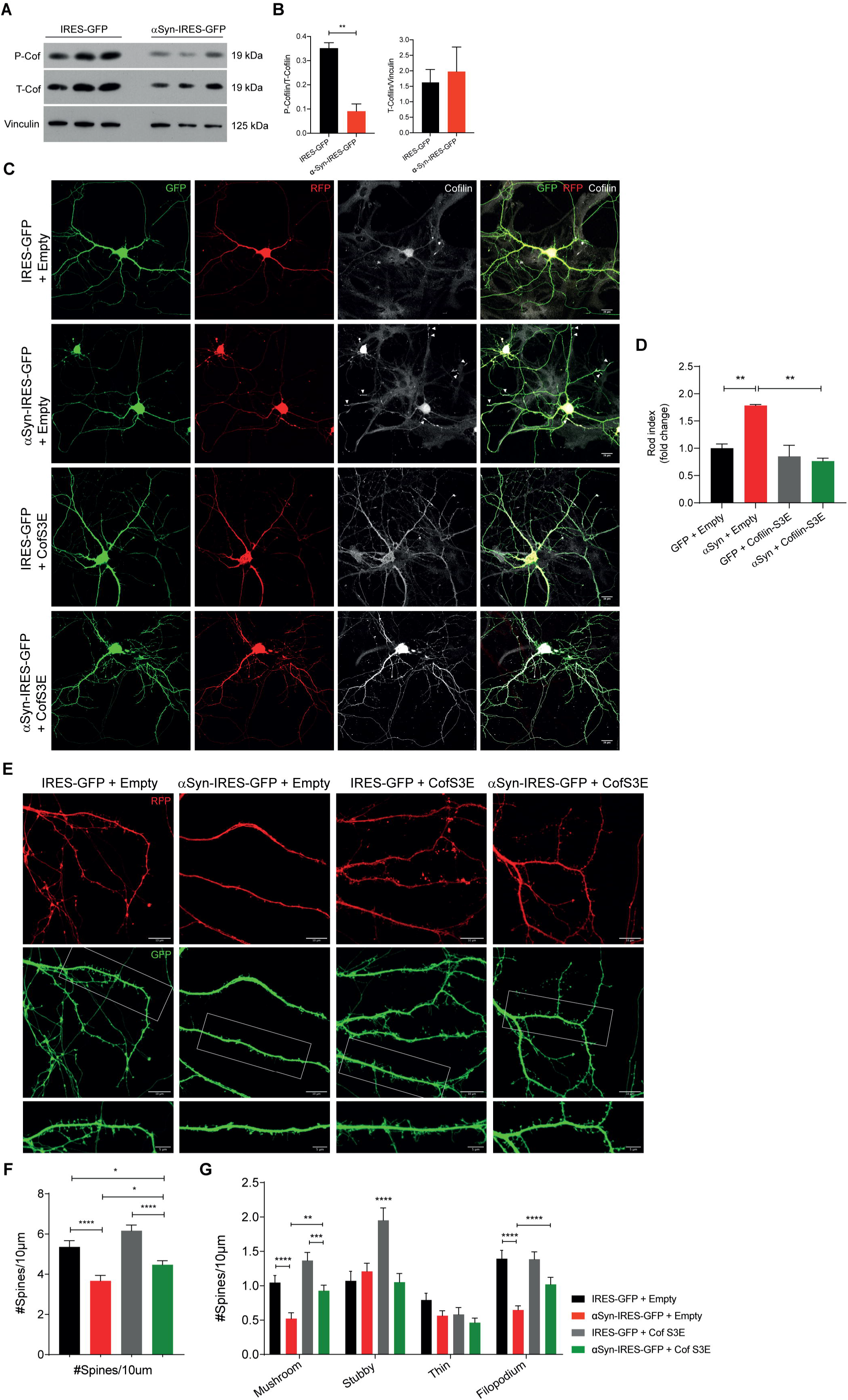
Cofilin pathology underlies α-Synuclein-induced rod formation and dendritic spine impairment. **(A, B)** Western blot analysis **(A)** and respective quantification by densitometry **(B)** of Cofilin pS3 and total Cofilin levels in DIV14 transduced hippocampal neurons. Vinculin was used as loading control. Data represent mean±SEM (n=3 replicates/treatment). **p<0.01 by Student’s *t* test. **(C)** Representative images of primary hippocampal neurons transfected at DIV12 with IRES-GFP or αSyn-IRES-GFP plasmids either with empty (pmRFP-N1) or Cofilin-S3E (pmRFP-N1-Cofilin-S3E) plasmids, and immunostained at DIV14 with Cofilin (white). Scale bar: 20μm. Arrowheads indicate cofilin-actin rod structures. **(D)** Rod index (fold change relative to control) relative to **C**. Data represent mean±SEM (n=2-3 replicates/treatment with 17-50 neurons/replicate). *p<0.05 by Student’s *t* test. **(E)** Representative images of primary hippocampal neurons transfected at DIV12 with IRES-GFP or WT-αSyn-IRES-GFP plasmids either with Empty (pmRFP-N1) or Cofilin-S3E (pmRFP-N1-Cofilin-S3E) plasmids, and visualized with GFP and RFP signals at DIV14. Scale bar: 10μm. Insets represent the respective zoom-ins noted by the white box. Scale bar: 5μm. **(F)** Dendritic spine density relative to **E**. Data represent mean±SEM (n=14-25 dendrites/treatment). *p<0.05, ****p<0.0001 by Student’s *t* test. **(G)** Dendritic spine density by morphology relative to **E**. Data represent mean±SEM (n=14-25 dendrites/treatment). **p<0.01, ***p<0.001, ****p<0.0001 by 2-way ANOVA with Sidak’s multiple comparison test.

**Figure 4.**
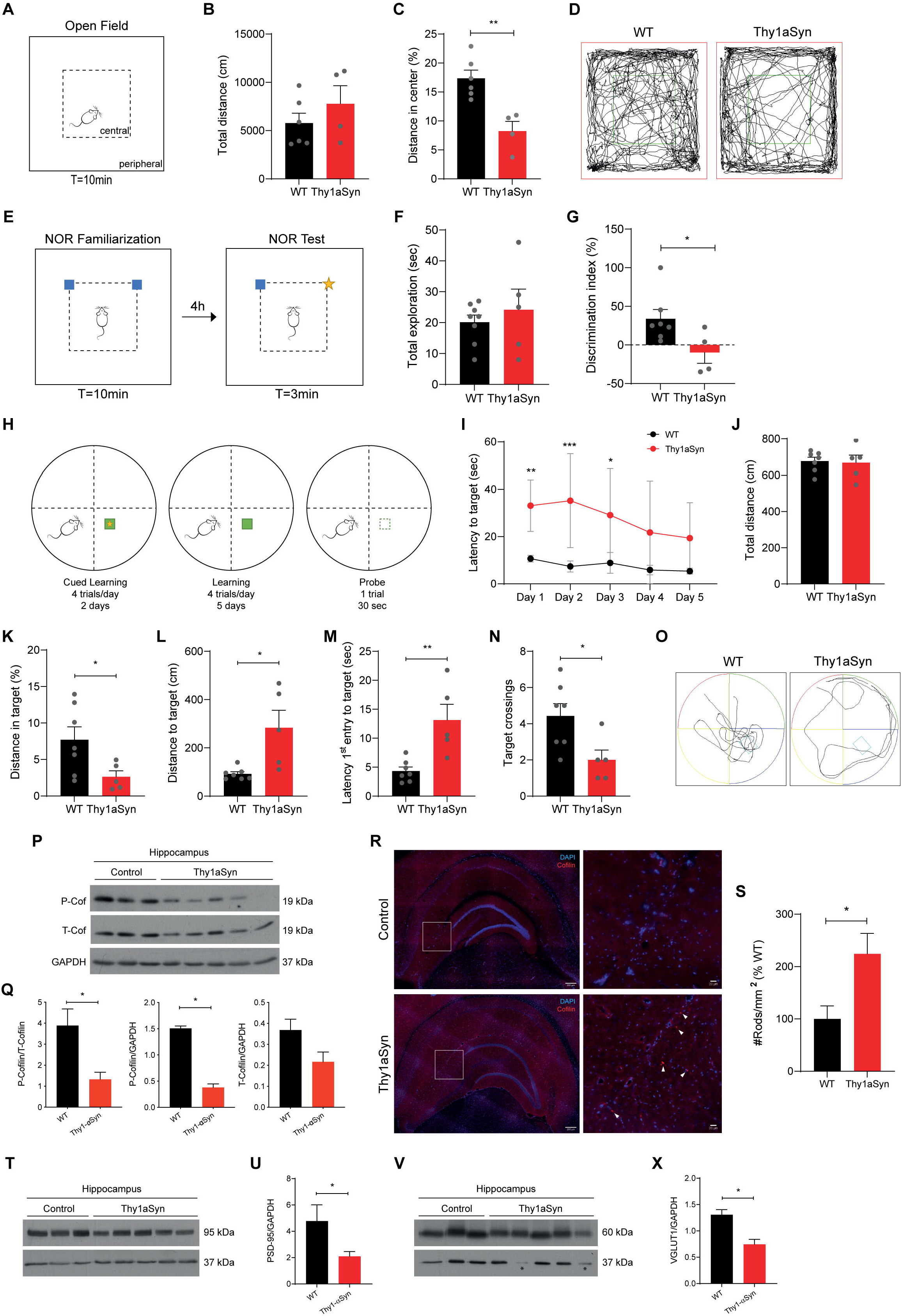
Cofilin pathology is observed in the Thy1-aSyn mice with cognitive impairment. **(A-D)** Open-field test in 6-month mice. **(A)** Schematic representation of the open field arena with the central area denoted in dashed line. **(B)** Total distanced traveled during the 10 min of the open field test. **(C)** Percentage of distance spent in the central area of the arena. Data represent mean±SEM (n=4-6 animals/condition). **p<0.01 by Student’s *t* test. **(D)** Representative tracks of WT and Thy1-aSyn mice on the open field test. **(E-G)** Novel object recognition (NOR) test in 6 months animals. **(E)** Schematic representation of the NOR test. **(F)** Total exploration time of the familiar and new objects in the NOR test. **(G)** Percentage of discrimination index in the NOR test. Data represent mean±SEM (n=4-7 animals/condition). *p<0.05 by Student’s *t* test. **(H-O)** Morris Water Maze (MWM) test in 6 months animals. **(H)** Schematic representation of the MWM test. **(I)** Latency to target in the learning phase. Data represent mean±SEM (n=5-7 animals/condition). *p<0.05, **p<0.01 by Student’s *t* test. **(J-O)** Probe trial with analyses of the total distance **(J)**, distance in target **(K)**, distance to target **(L)**, latency for the 1^st^ entry to target **(M)**, target crossings **(N)** and a representative track of the WT and Thy1-aSyn mice during the probe test **(O)**. Data represent mean±SEM (n=5-7 animals/condition). *p<0.05, **p<0.01 by Student’s *t* test. **(P, Q)** Western blot analysis **(P)** and respective quantification by densitometry **(Q)** of Cofilin pS3 and total Cofilin levels in hippocampus from Control and Thy1-aSyn mice. GAPDH was used as loading control. Data represent mean±SEM (n=3-5 animals/condition). *p<0.05 by Student’s *t* test. **(R)** Representative images of brain sections from Control and Thy1-aSyn mice with 6 months of age and stained for DAPI (blue) and Cofilin (red) and the respective insets in the hippocampal region. Scale bar: 200μm and 20μm for insets. Arrowheads indicate cofilin-actin rod structures. **(S)** Number of rods per mm^2^ (percentage relative to WT) in the hippocampus region relative to **R**. Data represent mean±SEM (n=4-5 animals/condition with 2-3 sections/animal). *p<0.05 by Student’s *t* test. **(T, U)** Western blot analysis **(T)** and respective quantification by densitometry **(U)** of PSD-95 levels in hippocampus from Control and Thy1-aSyn mice. GAPDH was used as loading control. Data represent mean±SEM (n=3-5 animals/condition). **(V, X)** Western blot analysis **(V)** and respective quantification by densitometry **(X)** of VGLUT1 levels in hippocampus from Control and Thy1-aSyn mice. GAPDH was used as loading control. For the purpose of quantification, the two animals with very low levels of GAPDH* were not considered. Data represent mean±SEM (n=3 animals/condition).

### Cofilin hippocampal pathology is observed in a mouse model of synucleinopathy with cognitive impairment

Our *in vitro* data revealed that αSyn overexpression leads to an hyperactivation of cofilin in primary cultures of hippocampal neurons which results in cofilin-actin rod formation and dendritic spine impairment.

We next aimed to determine whether αSyn-induced hippocampal cofilin pathology is recapitulated *in vivo*. We used a mouse model overexpressing human WT αSyn under the control of the neuronal Thy-1 promoter (Thy1-aSyn mice)^30^. This model recapitulates the αSyn levels observed in patients with multiplications of the *SNCA* gene and presents synaptic and memory impairments starting at early stages, similarly to what is seen in PD patients who develop dementia.

Using 6-month animals, an age where synaptic and cognitive dysfunction in Thy1-aSyn was characterized^27^, we initially confirmed the overexpression of human αSyn in the hippocampus as well as the presence of its pathologic-associated form αSyn pS129, by immunostaining and western blot (Figure S3A-C). We also validated cognitive impairment in the Thy1-aSyn mice. The open field test (OFT) was primarily performed to evaluate anxiety-like behaviors and locomotion and to habituate the animals to the arena used in the subsequent test (Figure 5A). While the total distance traveled in the open field arena was not different between control and Thy1-aSyn (Figure 5B and D), revealing no locomotor defects, we observed that the percentage of distance traveled in the central sector of the box was decreased in the Thy1-aSyn mice (Figure 5C and D), suggesting that Thy1-aSyn mice present an anxiety-like behavior as previously described^30^.

Next, we evaluated hippocampal-related cognitive functions namely non-spatial and spatial memories tested by Normal Object Recognition (NOR) and Morris Water Maze (MWM) tests, respectively. In NOR test (Figure 5E), the total time of exploration of the objects was not different between the two experimental groups (Figure 5F). However, whereas WT mice spent more time exploring the new object, the Thy1-aSyn mice did not distinguish between the familiar and new objects, as measured by the discrimination index (Figure 5G). These results validate the decreased recognition memory which was previously reported for the Thy1-aSyn mice^30^. In the MWM test (Figure 5H), Thy1-aSyn already presented defects in the learning phase showing increased latency to find the platform (Figure 5I). The WT control animals did not show a major improvement during the learning phase since pretraining (cued learning) reduced both the stress levels and the relative deficit in spatial learning expected in the first session in control mice (Figure S3D). In the probe test, although Thy1-aSyn mice travelled the same distance as WT littermates (Figure 5J and O), they spent less time on the target (Figure 5K and O), travelled an increased distance to reach the target (Figure 5L and O), had an increased latency in the first entry to target (Figure 5M and O) and made fewer target crossings (Figure 5N and O). Although the distance spent in the target quadrant and the mean distance to target were not significantly different, they had a tendency to decrease and increase, respectively, in the Thy1-aSyn mice (Figure S3E and F). Together, these results confirm that Thy1-aSyn mice show impaired learning and reference memory.

Having a model of validated hippocampal αSyn pathology and cognitive impairment, provided an excellent tool to validate *in vivo* our initial hypothesis that αSyn-induced cofilin pathology plays a role on hippocampal synaptic dysfunction and cognitive deficits in Lewy Body dementias. As such, we analyzed the activation state of cofilin in the hippocampus from the 6-months old Thy1-aSyn mice. Western blot analysis revealed decreased levels of cofilin pS3 in the Thy1-aSyn mice with no alterations in the total form of the protein, validating *in vivo* αSyn-induced activation of cofilin in the hippocampus (Figure 5P and Q). Moreover, cofilin activation was accompanied by a 2.2-fold increase in rod formation in the hippocampus of Thy1-aSyn mice (Figure 5R and S). We followed by analyzing whether cofilin activation was accompanied by synaptic dysfunction in Thy1-aSyn mice, by measuring the levels of the pre-synaptic protein VGLUT1 and the post-synaptic protein PSD-95 in hippocampal protein extracts. Western blot results showed decreased levels of both synaptic proteins in Thy1-aSyn mice when compared with WT controls (Figure 5T-X). Interestingly, the decreased levels of PSD-95 follow the decrease in dendritic spine density observed *in vitro*.

Our *in vivo* data shows a correlation between hippocampal cofilin pathology, synaptic defects and cognitive dysfunction in Lewy Body dementias but does not show a definitive cause and effect relationship. Further work will address the impact of modulating cofilin activity *in vivo* in rescuing synaptic impairment and cognitive deficits in Thy1-aSyn mice.

## Discussion

Cognitive dysfunction and dementia symptomatology critically affects functioning and quality of life of patients with Lewy Body dementias. These symptoms have been related to αSyn hippocampal pathology, as αSyn-containing inclusions are regularly detected in the hippocampus of patients with synucleinopathies, which present hippocampal atrophy and dysfunction as well as cognitive deficits^4^. Nevertheless, the specific contribution of the hippocampus needs further investigation in a proper model, being important to determine the pathologic consequences of αSyn accumulation in that brain region. The major findings of this work demonstrated both *in vitro* and *in vivo* that αSyn overexpression in the rodent hippocampus induces cofilin activation by dephosphorylation and cofilin-actin rods formation. In αSyn-overexpressing primary cultures of hippocampal neurons, cofilin activation partially mediated dendritic spine impairment. Importantly, and in accordance with our *in vitro* data, we correlated hippocampal cofilin pathology triggered by αSyn with synaptic dysfunction and cognitive impairment *in vivo* by using the Thy1-aSyn mice, a suitable model to study cognitive deficits in the context of Lewy Body dementias^31^.

Cofilin dysregulation has been previously associated to αSyn pathology in primary cultures of hippocampal neurons. Studies by Chieregatti et al reported an αSyn-induced activation of the actin signaling pathway Rac1/PAK2/LIMK/cofilin-1 *via* GRP78 in hippocampal neurons resulting on cofilin phosphorylation and inactivation, and consequent blockage of actin dynamics^32, 33^. Here we present an opposite observation with αSyn inducing cofilin activation in hippocampal neurons. This discrepancy might be related with distinct experimental conditions of αSyn exposure. While in the reported study there was an acute addition of 1µM recombinant αSyn to DIV14 neurons^32^, in our settings hippocampal neurons were transduced at DIV4 and analyzed at DIV14 where we observed a 3-fold increase in αSyn levels. Additionally, we validated cofilin activation *in vivo* in the hippocampus of Thy1-aSyn mice with overexpression of αSyn. These overexpression scenarios are more suitable for our propose of studying hippocampal αSyn pathology in the context of Lewy Body dementias as they recapitulate αSyn levels detected in patients with multiplications of the *SCNA* gene, which often develop cognitive deficits. Nevertheless, it would be interesting to further analyze the activation status of cofilin in additional mouse models and also focus on other cell types such as dopaminergic neurons, the ones mainly affected in typical motor PD, to determine whether αSyn-induced cofilin activation has neuronal subtype specificity.

Subsequently to cofilin activation induced by αSyn overexpression, we observed the formation of cofilin-actin rods in hippocampal neurons. Although not related to hippocampal pathology, cofilin-actin rod formation was previously reported in a *Drosophila* model of αSyn overexpression^34^. Neuronal rod formation was induced not only by αSyn overexpression, but also in response to the exogenous addition of αSyn soluble forms released from neurons and of αSyn PFFs to hippocampal neurons. Focusing on the data with mature DIV14 neurons, which reflect a more physiologic scenario, it might suggest that the intraneuronal accumulation of αSyn pS129 has a major contribution to rod formation, as shown in the experiments of long-term αSyn overexpression or αSyn PFFs addition to neurons (Figure 2B). Nevertheless, in those experiments, αSyn was present in the extracellular milieu, what combined with the observations showing that short-term addition of αSyn released from neurons or of αSyn PFFs resulted in rod formation, suggest an extracellular αSyn-interaction with a membrane receptor. In this respect, αSyn-induced rods was mediated by a PrP^C^-NOX pathway similarly to what was reported for Aβ^25^ and more recently for the HIV gp120 envelope protein^35^. Considering these evidences, we suggest that hippocampal rod formation seems to be the result of a combined cell-autonomous and non-cell autonomous mechanism which results from intraneuronal αSyn aggregation and αSyn spreading, respectively. αSyn spreading can lead to a signaling cascade at the cell membrane or to αSyn uptake and seeding of the endogenous protein. These conclusions were supported not only by our *in vitro* data but also by the *in vivo* data with the αSyn PFFs-injected mice and with the Thy1-aSyn mice.

Rod formation was shown to impair synaptic function in hippocampal neurons^11^, what was validated in our model by showing that cofilin pathology is involved in spine impairment induced by αSyn. Interestingly, PrP^C^ was previously described to interact with αSyn, mediating hippocampal synaptic impairment and cognitive deficits in PD^27^. Considering the inhibitory effect of PrP^C^knockdown on αSyn-induced rod formation, our data includes cofilin pathology as a new intermediate on the mechanism of αSyn-PrP^C^induction of synaptic and cognitive dysfunction.

Overall, our results strengthen the link between αSyn-induced hippocampal synaptic dysfunction and cognitive deficits in Lewy Body dementias and identify cofilin as a novel pathologic player. Future work will address the effect of targeting cofilin pathology in the Thy1-aSyn mice, to validate our hypothesis that cofilin dysregulation is responsible for synaptic dysfunction in the context of dementia in synucleinopathies. Current treatments for Lewy Body dementias include mainly the use of cholinesterase inhibitors, as cholinergic deficit in patients with dementia was reported. However, efficacy and safety of cholinesterase inhibitors needs further analysis in larger clinical trials^36, 37^. Moreover, PD treatments mainly focus on increasing neurotransmitter signaling and it is important to develop new pharmacological approaches to prevent or reverse synapse loss and neuronal connectivity. As such, the development of new therapies targeting dementia in the context of synucleinopathies is imperative, and our future goal of targeting cofilin pathology, which impacts on neuronal dysfunction, constitutes a promising innovative strategy impacting not only on Lewy Body dementias but also in additional disorders, namely AD and HIV-associated Neurocognitive Disorder (HAND), where rods were reported.

## Materials and Methods

### Animals

Animal procedures were performed in accordance with national and European rules. The protocols described in this work have been approved by the IBMC Ethical Committee and by the Portuguese Veterinarian Board and by the Institutional Animal Care and Use Committee of Colorado State University (protocols KP1023 and KP1412). Transgenic mice overexpressing human αSyn under the neuronal Thy-1 promoter (Thy1-aSyn mice, Line 61 developed on a C57Bl6/DBA2 background)^38^ were kindly provided by University of California San Diego. The colony background was maintained by breeding mutant females with WT C57BL/6-DBA/2 males. Since the transgene insertion was on the X chromosome and there is random inactivation of the X chromosome, only male littermates were used in the experiments.

At 6 months of age animals were used in behavioral tests, sacrificed and their brains collected. For immunostaining analysis, mice were perfused with PBS for 5 min followed by 4% paraformaldehyde (PFA, pH 7.4) in PBS (40ml). Brains were incubated in 4% PFA for 24h and then in 30% sucrose. For western blot analysis mice were perfused with PBS for 5 min, the hippocampus dissected and quickly frozen in dry ice.

For αSyn PFFs stereotactic injection, a microinjection syringe was inserted to target the substantia nigra pars compacta bilaterally (anterior–posterior, +/-3.0, medio-lateral, + 1.5, dorso-ventral, - 4.6 from bregma). Each injection delivered 10 µl of 5 μg/μL PFFs or, for controls, 10 µL of PBS. Post-surgical incisional pain was treated with bupivacaine and buprenorphine.

### Behavioral tests

#### Open-field (OF)

WT and Thy1-aSyn mice were subjected to the OF test during the dark period to evaluate activity levels. The system comprised an empty opaque squared arena (40×40 cm). Individually, mice were placed in the center of the arena and left to explore the area for 10 min. The box was cleaned between animals to eliminate odor or traces from the previous animal. Mice behaviors, as total distance travelled and activity in the central and peripheric areas, were recorded and posteriorly analyzed by the software The Smart v3.0, Panlab, Barcelona, Spain.

#### Novel object recognition (NOR)

Animals that have been previously tested in the OF (habituation) were subsequently subjected to the NOR test to evaluate memory. In the same arena used in the OF, animals were exposed to two identical objects for 10 min for familiarization. In the test phase, four hours later, one of the familiar objects was replaced by a new object from the same material, weight and height, but with a different color and shape, and the animal was allowed to explore for 3 min. Exploration was considered when the mouse’s nose touched the object or when the nose is directed to the object from a distance less than 2 cm. Mouse behaviors were recorded and analyzed by the software The Observer XT v7.0, Noldus, Netherlands. The discrimination index was calculated by DI = (*T*_N_ − *T*_F_)/(*T*_N_ + *T*_F_) in which *T*_N_ is the time exploring the novel object and *T*_F_ is the time spent exploring the familiar object. Animals with a total exploration time (novel + familiar) less than 10 seconds were excluded from the analysis of the discrimination index^39^.

#### Morris water maze (MWM)

Spatial learning was assessed by the hidden-platform Morris water maze test. A circular pool (diameter 111cm) was used and filled with water at 21±1ºC. The pool was theoretically divided in four quadrants and eight start positions were defined at equal distance to the center. An escape platform (10×10cm) was placed 0.5cm below the water line. In first two days, during the cued leaning, mice were trained to find the hidden platform which had a visual clue. Animals were subjected to four swimming trials which had a different start and goal position. In the next 5 days, during the learning phase, mice were trained to find the hidden platform which is in the same location during the entire learning phase. Every day mice were subjected to four swimming trials, each trial starting at one of the four different pool locations, and the latency to find the platform was scored. If mice failed to find the platform within 1 min they were guided to the platform. In either case, mice were allowed to stay on the platform for 15 sec. On day 8, in the probe day, the platform was removed from the pool and the mice were allowed to swim for 30 sec. Swimming tracks were recorded and analysis of the total distance, distance to target, latency for the 1^st^ entry to target and target crossings were obtained with The Smart v3.0, Panlab, Barcelona, Spain.

### Primary hippocampal neuron cultures

The hippocampus was dissected from E18 rat or E16.5 mouse embryos, digested with 0.06% trypsin (Sigma-Aldrich, T4799) in Hanks’ balanced salt solution (HBSS, Sigma, H9394) for 15 min at 37^º^C. Following digestion, neurons were dissociated by gentle trituration and resuspended in neurobasal medium (Invitrogen, 21103049) supplemented with 2% N21 (R&D Systems, AR008), 1% penicillin/streptomycin (ThermoFisher Scientific, 15140-122) and 0.25mM L-Glutamine (Lonza, 17-605E). Cells were then counted and platted at a density of 15,000 cells/coverslip in a 24-well plate for immunostaining analysis, or at a density of 200,000 cells/well in a 6-well plate for western blot analysis. Coverslips and plates were precoated with 20 µg/mL poly-D-lysine (Sigma, P0899). For immunostaining cells were fixed with 4% PFA in cytoskeleton preservation buffer (10mM 2-(N-morpholino) ethanesulfonic acid (MES, Sigma-Aldrich, M3671) pH 6.1; 3mM MgCl_2_ (Merk, 1.05833.0250); 138mM KCl (Merk, 529552); 2mM ethylene glycol-bis (β-aminoethyl ether)-N,N,N’,N’-tetraacetic acid (EGTA, Sigma-Aldrich, E8145); 0.32M sucrose (Merk, 1.07651.1000)), for 30 min at room temperature (RT). For western blot, cells were harvested and lysed in 0.3% triton X-100 (Sigma-Aldrich, T9284), 1x protease inhibitor Cocktail (100x, GE Healthcare, 80-6501-23) and 1mM sodium orthovanadate (Sigma-Aldrich, S6508).

### Plasmids and viral vectors

IRES-GFP and WT αSyn-IRES-GFP lentiviral plasmids were previously described^40^. Briefly, full-length human WT αSyn c-DNA was subcloned into the pWPI vector (second generation bicistronic lentiviral vector, Tronolab, Switzerland), under the chicken/β-actin (CBA) promoter. A pWPI vector containing only IRES-GFP was used as control. pmRFP-N1 and pmRFP-N1-Cofilin-S3E plasmids were previously described^41^. Briefly, psuedo-phosporylated human cofilin-1 (S3E) was cloned into the pmRFP-N1 backbone vector, under the CMV promoter. An empty pmRFP-N1 vector was used as control.

### Lentiviruses production and titration

Lentivirus production was performed as previously described^42^. Briefly, HEK293T cells were transfected, using Lipofectamine 2000 (ThermoFisher Scientific, 11668030), with the DNA complexes containing the plasmid of interest and the packaging plasmids (psPAX2 and VSV-G), for 5h at 37^º^C/5%CO_2_. After the incubation, medium was replaced with DMEM (VWR, 733-1695) supplemented with 10% FBS (Biowest, BWSTS181BH-500) and 1% P/S. After 48h, the lentivirus-containing supernatants were recovered, centrifuged for 10 min at 500g and filtered using a 0.45um filter (Enzifarma). The filtered supernatants were concentrated using a centricon (GE Healthcare Life Sciences), aliquoted and stored at -80^º^C. For virus titration, HEK293T cells were infected with different volumes of lentivirus. After 3 days, cells were resuspended in PBS and the total number of transduced cells was analyzed by Flow Cytometry using FACS Accuri (BD Biosciences). The lentiviral transfection units (TU) per μL were determined by the following equation: TU/μL= (number of plated cells x % of infected cells (GFP positive)) / volume of viral particles added (μL).

### αSyn monomers, oligomers and pre-formed fibrils (PFFs)

αSyn monomers and oligomers were prepared from recombinant human αSyn as previously described^43^. Prepared species were aliquoted and stored at -80°C and thawed in ice immediately before use. For PFF preparation, human αSyn monomer protein was purchased from Proteos (RP-003). PFF were prepared according to the protocol stablished by Michael J Fox Foundation for Parkinson’s Research^44^. Briefly, αSyn monomers were diluted to 5 mg/ml into 0.01M phosphate buffered saline (PBS) containing 0.03% sodium azide to prevent bacterial growth. Monomers were shaken for 7 days at 1000 rpm and 37°C in an orbital shaker to induce the formation of fibrils (PFF). Single-use aliquots were rapidly frozen and stored at -80 °C. Immediately before use, frozen aliquots were thawed, diluted in PBS to 0.1 mg/mL and bath sonicated at room temperature for 5 minutes.

### Hippocampal neurons transduction

DIV 3 hippocampal neurons were treated with 1μM of (+)-MK-801 hydrogen maleate (Sigma, M107) for 30 min at 37^º^C to reduce spontaneous rod formation. At DIV4, neurons were infected either with WT αSyn-IRES-GFP or IRES-GFP lentiviruses (15.000TU/condition). At DIV7 or DIV14 culture medium was recovered and cells fixed as described in the previous section.

### Hippocampal neurons transfection

DIV 3 hippocampal neurons were treated with 1μM of (+)-MK-801 hydrogen maleate (Sigma, M107) for 30 min at 37^º^C to reduce spontaneous rod formation. At DIV12 neurons were transfected using the calcium phosphate co-precipitation method with the constructs: αSyn-IRES-GFP, IRES-GFP, pmRFP-N1-Cofilin-S3E, and pmRFP-N1. Briefly, a maximum amount of 2μg of DNA (single or mixture) was diluted in Tris-EDTA (TE) pH7.3 and mixed with HEPES calcium chloride (2.5M CaCl2 in 10mM of HEPES pH7.2). This mixture was added to 2x HEBS (270mM NaCl, 10mM KCl, 1.4mM Na2HPO4, 11mM Dextrose, 42mM HEPES pH7.2) and the precipitate was allowed to develop during 30 min at RT in the dark, with gentle mixing every 5 min. Culture medium was removed and saved while neurons were incubated with plain neurobasal medium and the precipitates were added dropwise to each well. Precipitates were incubated with cells for 45 min at 37^º^C/5%CO_2_. Precipitate solution was then removed and neurons washed with acidic neurobasal medium (equilibrated at 10%CO_2_) for 20 min at 37^º^C/5%CO_2_. Lastly, the medium was replaced with the saved supplemented medium. Neurons were analyzed at DIV14, 48h after transfection.

### Immunocytochemistry

For cofilin-actin rods staining, neurons were permeabilized with 100% methanol at -20^º^C for 3 min at RT and blocked with 2.5% normal serum from donkey (Jackson ImmunoReasearch, 017-000-121) or goat (Sigma, 19H092) in 1%BSA/PBS for 1h at RT. Followed incubation with primary antibodies: rabbit anti-T-Cofilin 1:2000 (Bamburg lab, 1439 or Cell Signalling, 5175) and mouse anti-β3-tubulin 1:2000 (Promega, G7121) diluted in 1%BSA/PBS and incubated overnight at 4^º^C. After washing, cells were incubated with secondary antibodies: donkey anti-mouse-Alexa Fluor 488 1:1000 (Invitrogen, A21202) or donkey anti-mouse-Alexa Fluor 647 1:1000 (Invitrogen, A31571) and donkey anti-rabbit–Alexa Fluor 568 1:1000 (Invitrogen, A10042) diluted in 1%BSA/PBS. Coverslips were mounted in Fluoromount-G (SouthernBiotech, 0100-01) or ProLong Diamond Antifade (ThermoFisher P36961). For αSyn staining, neurons were permeabilized with 2.5% triton X-100 in PBS for 20 min at RT and blocked with 5% normal donkey serum in 1%BSA/PBS for 1h at RT. Subsequently, neurons were incubated with primary antibodies: mouse anti-αSynuclein 1:1000 (BD Biosciences, 610787) and rabbit anti α-Syn pS129 1:1000 (Abcam, ab51253) diluted in 1%BSA/PBS and incubated overnight at 4^º^C. After washing and incubation with secondary antibodies: donkey anti-rabbit–Alexa Fluor 568 1:1000 (Invitrogen, A10042) and donkey anti-mouse-Alexa Fluor 647 1:1000 (Invitrogen, A31571) diluted in 1%BSA/PBS, coverslips were mounted in Fluoromount-G.

### Immunohistochemistry

Brain tissues were embedded in Optimum Cutting Temperature (OCT) compound (ThermoFisher Scientific), frozen and sectioned coronally (Cryostat Leica CM3050S) at 30μm. For cofilin-actin rods staining sections were permeabilized with 100% methanol at -20^º^C for 5 min at RT and blocked with 5% normal donkey serum in PBS for 1h at RT, followed by incubation overnight at 4^º^C with primary antibodies: rabbit anti-T-Cofilin 1:1000 and 1:1000; and subsequent washing and incubation with secondary antibodies: donkey anti-mouse-Alexa Fluor 488 1:500 and donkey anti-rabbit–Alexa Fluor 568 1:500 diluted in 1%BSA/PBS. Brain sections were then washed and rinsed in 70% ethanol and incubated with Sudan Black in 70% ethanol for 10 min at RT. Sections were washed, incubated with DAPI (Bio-Rad, 1351303) for 10 min and mounted in ibidi mounting medium (ibidi, 50001).

For staining of α-Syn pS129, sections were washed with 0.3% triton X-100 in PBS and blocked with 1% normal donkey serum in 0.3% triton X-100 in PBS for 1h at RT. Followed by incubation with primary antibody: rabbit anti-α-Syn pS1291:5000 (Abcam, ab51253) diluted in blocking buffer and incubated 2 days at 4^º^C. Following washing and incubation with secondary antibodies: donkey anti-rabbit–Alexa Fluor 568 1:1000, sections were washed, incubated with DAPI for 10 min and mounted in ibidi mounting medium.

### Western blot

Frozen brains were incubated with RIPA buffer (1% Triton X-100, 0.1% SDS, 140mM NaCl, 1x TE pH 8, 1x protease inhibitor Cocktail and 1mM Sodium orthovanadate), sonicated (2×10 cycles, Output Power 50 Watts, Branson sonifier 250) and cleared by centrifugation at 15000rpm for 5 min at 4^º^C. Neuronal cell lysates were sonicated (2×10 cycles, Output Power 50 Watts, Branson sonifier 250) and cleared by centrifugation at 15000rpm for 5 min at 4^º^C. 25ug or 5ug of protein extracts were separated under denaturating conditions in 12% SDS-PAGE gels and transferred to nitrocellulose membranes (0.45μm GE HealthCare) for 2 hours, using a semi-dry transfer system (CBS scientific EBU-4000). Membranes were blocked with 5% milk (Sigma-Aldrich) in TBS-T or 5%BSA (NZYTech) in TBS-T for 1 h at RT. Membranes were probed overnight at 4^º^C with the following primary antibodies: mouse anti-αSynuclein 1:1000 (BD Biosciences, 610787), rabbit anti-α-Syn pS129 1:500 (Abcam, ab51253), rabbit anti-T-Cofilin 1:1000 (Bamburg lab, 1439 or Cell Signaling, 5175), mouse anti-T-Cofilin 1:500 (Abcam, ab54532), rabbit anti-P-Cofilin Ser3 1:1000 (Cell Signaling, 3311), mouse anti-PSD-95 1:2000 (ThermoFisher Scientific, MA1-046), rabbit anti-VGLUT1 1:5000 (Synaptic Systems, 135303), rabbit anti-Vinculin 3:10000 (ThermoFisher Scientific, 700062) and mouse anti-GAPDH 1:1000 (Santa Cruz, sc-166574) diluted in 5%milk/TBS-T or 5%BSA. After washing the membrane was incubated for 1h at RT with secondary antibodies: anti-mouse IgG-HRP 1:10000 (Jackson Research, 115-035-003) or anti-rabbit IgG-HRP 1:10000 (Jackson Research, 111-035-003) diluted in 5% milk/TBS-T. Immunodetection was performed by chemiluminescence using ECL (Millipore, WBLUR0500). Quantitative analyses were performed with the Quantity One software or Fiji software.

### Dot blot

Supernatants collected from transduced hippocampal neurons were centrifuged at 4000g for 5 min at 4^º^C to remove cell debris and then applied to a nitrocellulose membrane using FisherbrandTM Dot Blot Hybridisation Manifold System according to the manufacturer’s recommended protocol. Membranes were allowed to dry and then hydrated and blocked for 1h at RT with 5%milk/TBS-T and incubated overnight at 4^º^C with the primary antibody mouse monoclonal anti-α-Syn 1:1000 (BD Biosciences, 610787) in 5%milk/TBS-T. After washing, membranes were incubated 1h at RT with secondary antibody anti-mouse IgG-HRP diluted in 5% milk in TBS-T. Immunodetection was performed by chemiluminescence using ECL reagent.

### Imaging and quantifications

Hippocampal neurons cultured for 7 days and immunostained for cofilin were assessed for the presence of cofilin-actin rods in an upright epifluorescence microscope (Zeiss Axio Imager Z1, Carl Zeiss) at 40x magnification. The percentage of neurons with cofilin-actin rods was plotted. Hippocampal neurons cultured for 14 days and immunostained for cofilin were imaged in an automated fluorescence widefield high-content screening microscope (IN Cell Analyzer 2000, GE Healthcare) at 40x magnification. Images were analyzed using Fiji software and the ratio between the number of rods and the total number of neurons analyzed was calculated and plotted as Rod index. Imaging of cofilin in DIV7 PFF-induced rods was performed on a Keyence Fluorescence Microscope with a 20x objective.

For spine density quantification, hippocampal neurons cultured for 14 days and expressing GFP, were imaged in a laser scanning Confocal Microscope Leica SP5 AOBS SE, using the 63x oil objective. Dendritic length and spine number and morphology were quantified using the semi-automatic NeuronStudio software. Dendritic spine morphology was defined as mushroom spines (small neck and large head), thin spines (long neck and small head), stubby spines (head without a defined neck) and filopodium spines (long neck without defined head).

Brain sections stained for cofilin were imaged in an automated fluorescence widefield high-content screening microscope (IN Cell Analyzer 2000, GE Healthcare) at 20x magnification. Images were stitched and the brain regions of interest analyzed using Fiji software and the results plotted as the number of rods per area.

### Statistical analysis

All measurements were performed with the researcher blinded to the experimental condition when possible. Data are shown as mean ±SEM. Statistical significance was determined using the GraphPad Prism Software version 8 and the most appropriate statistic test being significance determined by *p<0.05, **p<0.01, **p<0.001 and ****p<0.0001. Statistic test and sample sizes are indicated in each figure legend.

## Acknowledgements

We acknowledge University of California San Diego for providing the Thy1-aSyn mice. We thank the i3S Animal facility, Genotyping, Cell Culture, Histology and Electron Microscopy, Advanced Light Microscopy and BioSciences Screening (PPBI-POCI-01-0145-FEDER-022122) Facilities.

This work was supported by: FEDER - Fundo Europeu de Desenvolvimento Regional funds through the COMPETE 2020 – Operacional Programme for Competitiveness and Internationalisation (POCI), Portugal 2020, by Portuguese funds through FCT - Fundação para a Ciência e a Tecnologia/Ministério da Ciência, Tecnologia e Ensino Superior in the framework of the project POCI-01-0145-FEDER-028336 (PTDC/MED-NEU/28336/2017); R&D@PhD from Luso-American Development Foundation (FLAD); FLAD Healthcare 2020; and Programme for Cooperation in Science between Portugal and Germany 2018/2019 (FCT/DAAD). Research was also supported by generous gifts to the Colorado State University Development Fund (JRB) and by the National Institutes on Aging of the National Institutes of Health under award numbers R01AG049668 and R43AG071064 (JRB). The content is solely the responsibility of the authors and does not necessarily represent the official views of the NIH. TFO is supported by the Deutsche Forschungsgemeinschaft (DFG, German Research Foundation) under Germany’s Excellence Strategy - EXC 2067/1-390729940) and by SFB1286 (Project B8). MIOS is a FCT fellow (SFRH/BD/118728/2016). MAL is a FCT Investigator (IF/00902/2015).

