## Supplementary Material for "Cofilin pathology is a new player on α-synuclein-induced spine impairment in models of hippocampal synucleinopathy"

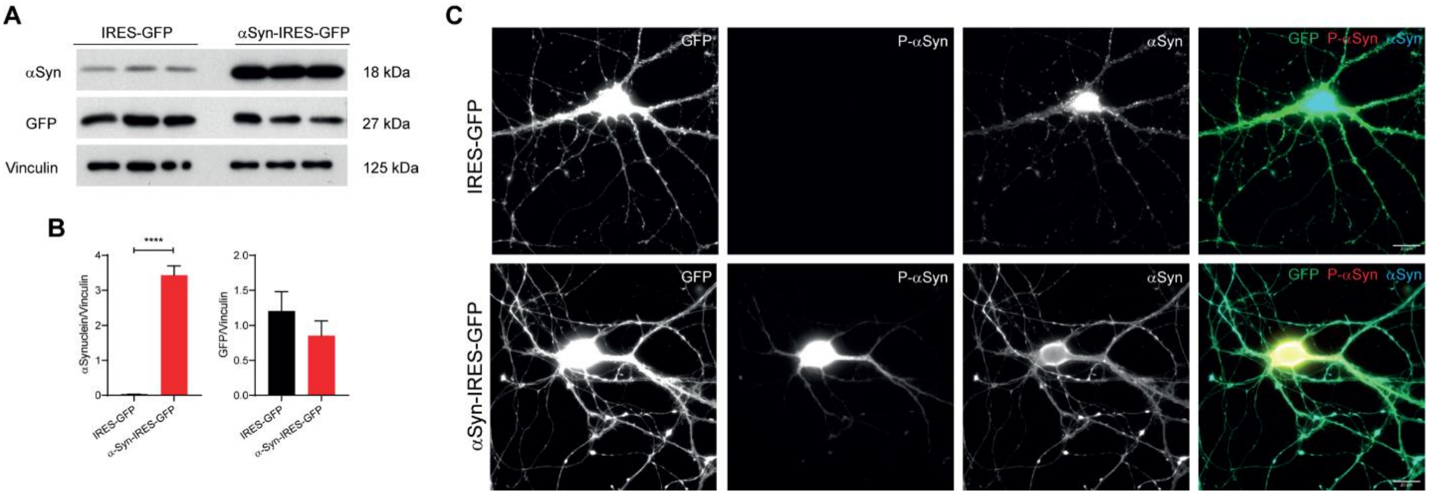


***Supplementary Figure S1 - Overexpression of αSyn in mature hippocampal neurons.*** **(A, B)** Western blot analysis **(A)** and respective quantification by densitometry **(B)** of α-Syn and GFP levels in DIV14 transduced hippocampal neurons. Vinculin was used as loading control. Data represent mean±SEM (n=3 replicates/treatment). ****$p<0.0001$ by Student's *t* test. **(C)** Representative images of primary hippocampal neurons infected at DIV4 with IRES-GFP or WT-αSyn-IRES-GFP lentivirus and immunostained at DIV14 with α-Syn pS129 (red) and α-Syn (blue). Scale bar: 20μm.

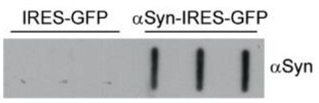


***Supplementary Figure S2 - αSyn is released from hippocampal neurons in culture.*** Dot blot analysis of αSyn levels in the supernatants from IRES-GFP or WT-αSyn-IRES-GFP transduced neurons at DIV7.

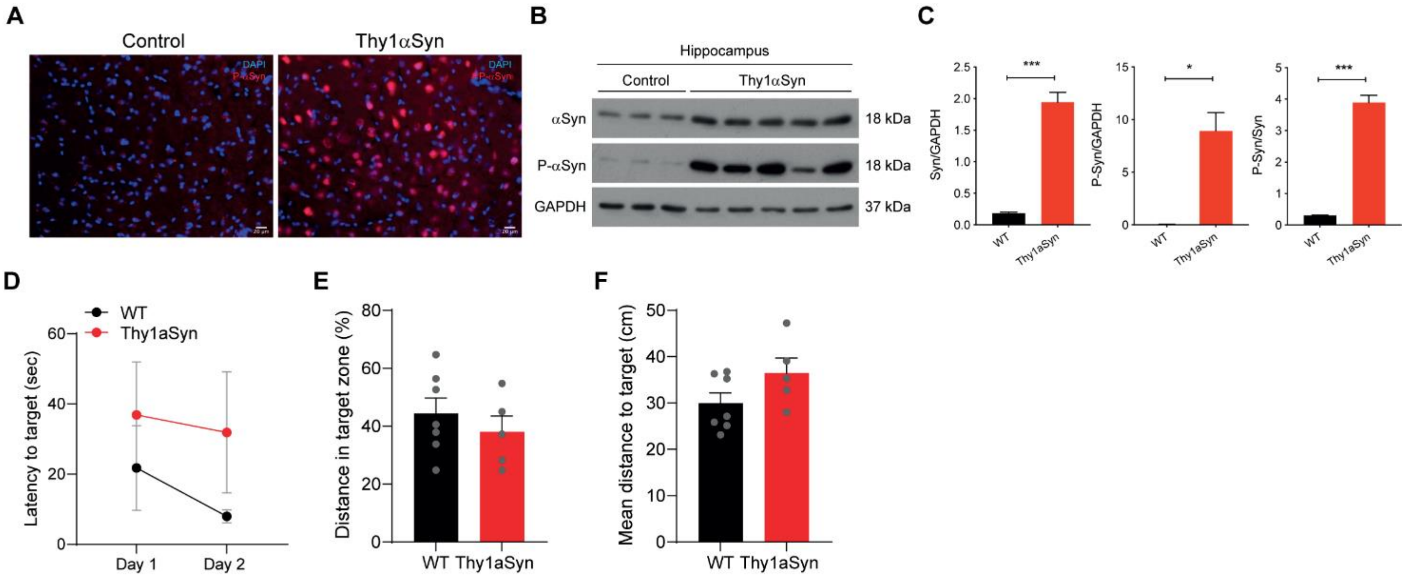


***Supplementary Figure S3 - Thy1-aSyn mice show increased αSynuclein phosphoS129 in hippocampus and cognitive defects.*** **(A)** Representative images of brain sections from 6-month Control and Thy1-aSyn mice stained for DAPI (blue) and α-Syn pS129 (red). Scale bar: 20μm. **(B, C)** Western blot analysis **(B)** and respective quantification by densitometry **(C)** of α-Syn and α-Syn pS129 levels in hippocampus from Control and Thy1-aSyn mice. GAPDH was used as loading control. Data represent mean±SEM (n=3-5 animals/condition). *p<0.05, ***p<0.001 by Student's *t* test. **(D)** Latency to target during cued learning in MWM. **(E)** Distance in target zone in MWM. **(F)** Mean distance to target in MWM. Data represent mean±SEM (n=5-7 animals/condition).
